# Dark adaptation of cone photoreceptor responses is revealed by optoretinography

**DOI:** 10.64898/2026.04.28.721480

**Authors:** Yao Cai, Anqi Zhang, Maciej M. Bartuzel, Reddikumar Maddipatla, Robert J. Zawadzki, Ravi S. Jonnal

## Abstract

Dark adaptation is the essential process that restores visual sensitivity following exposure to bright light, yet the underlying mechanisms remain incompletely understood. Here, we propose a method for assessing dark adaptation in cones using optoretinography (ORG) based on adaptive optics optical coherence tomography (AO-OCT). ORG quantifies cone functional response by monitoring nano-scale changes in the cone’s outer segment occurring over hundreds of milliseconds after visible stimulation. This method consists of sequential measurements of stimulus-evoked cone responses over the course of minutes of dark adaptation. Each response captures optical path length changes in single photoreceptor outer segments over milliseconds during a multi-minute recovery period following a strong photopigment bleach. We parameterized cone ORG responses and proposed an exponential model linking ORG dynamics to pigment regeneration. Parameters of the ORG response exhibited exponential decay behavior during dark adaptation, and were thus fit with exponential functions and quantified by the resulting decay parameter ***τ***. Parameters capturing the amplitude of the ORG responses recovered more slowly than those capturing temporal dynamics of the responses. This difference is consistent with distinct contributions from photopigment regeneration and downstream phototransduction processes. Recovery speed varied by two-to threefold among three normal subjects, suggesting substantial inter-subject physiological diversity. Processes within the cone, including pigment regeneration, are thought to underlie the gains in photopic visual sensitivity that occur in the dark. These findings highlight ORG as an objective and sensitive assay of those cellular mechanisms. While the ORG itself has shown promise as a biomarker of the health of the photoreceptor response to light, the results of this study show that it may also be useful for probing the health of the intra- and intercellular homeostatic mechanisms that support it.

## 1 Introduction

The vertebrate retina possesses the remarkable ability to operate under a broad range of light intensities spanning more than nine log units, a capability partly attributable to the adaptation of rod and cone photoreceptors [1, 2]. For example, when we go from a room where we are exposed to bright light to a dimly lit room where we cannot see at all, it takes some time for our eyes to adjust to this dark environment. In fact, the brighter the light in the first room, the longer it takes to adjust our vision in the darker room. Dark adaptation (DA) is the essential process by which these photoreceptors recover visual sensitivity in darkness following exposure to intense or prolonged illumination. This recovery is primarily driven by the regeneration of photopigments [3–6], though other mechanisms are thought to play a role both in cones and rods [7–11].

Upon exposure to light, photopigment molecules absorb photons and the chromophore 11-*cis* retinal is thereby isomerized to its all-*trans* form, losing its color, in what is known as bleaching of the photopigment (photobleaching). This process sets off the phototransduction cascade, causing the hyperpolarization of photoreceptor membrane [1, 12]. To restore light sensitivity, all-trans-retinal is released from opsin and recycled back to 11-cis-retinal in the retinal pigment epithelium (RPE). The 11-cis-retinal returns to the rod or cone outer segments and combines with opsin to regenerate the photopigment. This process, known as the visual cycle, forms the physiological basis for the recovery of retinal sensitivity during dark adaptation [4, 11]. Additionally, there is a ‘cone-specific’ visual cycle, allowing cone outer segments access to a separate pool of chromophore located within Müller cells, potentially contributing to a faster cone-driven recovery [2, 4, 5].

Dark adaptation depends in part on the biochemical recycling of all-trans-retinol back to 11-cis-retinal, a process which occurs across the interface of photoreceptor outer segments and the retinal pigment epithelium. Furthermore, DA depends on the supply of nutrients, including vitamin A and oxygen, which must traverse Bruch’s membrane (BrM) from the choriocapillaris. Thus, pathologic changes to any of these cell layers or intercellular spaces have the potential to impact dark adaptation. Photoreceptor sensitivity recovery is known to be slowed, and dark adaptation generally impaired, in patients with diseases affecting photoreceptor outer segments, RPE, Bruch’s membrane or choriocapillaris, including age-related macular degeneration (AMD) [4, 13]. Slower cone and rod dark adaptation has also been documented in eyes with age-related maculopathy [14, 15]. These functional deficits are particularly clinically significant because they may precede visible structural changes or any significant decline in visual acuity [4, 16].

Historically, three primary modalities have been developed for DA measurement: psychophysics, electrophysiology, and optical densitometry.

Psychophysical methods assess visual sensitivity by eliciting subject responses to light stimuli verbally or by pushing a button. These methods reveals a biphasic recovery curve representing rapid cone adaptation followed by slower rod-mediated recovery [5–7, 17, 18]. Key parameters derived from the resulting dark adaptation curve include absolute sensitivity thresholds for cones and rods, recovery rates, and time to cone-rod break (CRB) [4]. In clinical settings, dark adaptometers like the AdaptDx (OpZira, Inc.) utilize the rod-intercept time (RIT), defined as the time required for sensitivity to recover by 3 log units, as a standardized, reliable metric for quantifying rod function [16, 19]. However, these tests are inherently subjective, depend on patient cognitive health and cooperation, fixation stability, controlled pupil size, and ‘floor effects’ in severe disease, where sensitivity fails to reach measurable criteria [4].

Retinal densitometry optically measures the concentration and regeneration kinetics of photopigments in the living eye by quantifying the density of light reflected from the fundus [17, 20], but pigment depletion alone cannot account for the magnitude of psychophysical sensitivity loss observed during dark adaptation [2, 7, 21, 22].

Additionally, electrophysiological methods, primarily the electroretinogram (ERG), provide an objective measure of retinal dark adaptation by recording the electrical currents of photoreceptors (a-wave) and bipolar cells (b-wave). Previous studies found that rod circulating current was undetectable in the first 5 min and took 30 min to recover to dark-adapted levels after full bleach, due to the persistent effect of ‘equivalent background’ generated by unregenerated opsin [1, 7, 11, 23], though an early ‘fast adaptation’ mechanism mediated by phototransduction deactivation has been reported [24]. In contrast, cone current recovery occurs within milliseconds following similar bleaches, approximately 50,000-fold faster than rod recovery, indicating that bleaching products do not substantially suppress the circulating current in cones in the manner observed for rods. Importantly, the proportion of unbleached photopigment can be estimated by cone-driven a-wave amplitude, followed by a rate-limited recovery after bleaching[2, 25, 26]. A limitation of this approach is the potential for post-receptoral contamination in the cone-driven a-wave, arising largely from bipolar cell activity [15, 26], though this can be mitigated by quantifying a-waves very soon after the flash to avoid significant post-receptoral intrusion [2, 27]. Also, standard full-field ERG represents a massed response of the entire retina, lacking the spatial resolution to interrogate individual cells or localized pathology.

Despite significant progress linking pigment regeneration with electrophysiological and psychophysical recovery, fundamental questions remain. The precise mechanisms underlying the elevation of psychophysical threshold caused by light exposure and the contributions of processes occurring at different levels of the retina are still to be fully elucidated. Specifically, the role of proteins involved in phototransduction (e.g., transducin and arrestin) in modulating post-bleach sensitivity is not fully understood. Furthermore, the relative contributions of the retinal pigment epithelium versus Müller cell pathways to human cone pigment regeneration are unknown [2]. This uncertainty extends to the blue cone system, which exhibits slower regeneration than red or green cone systems, though the causes for the differences remain unclear [18]. Beyond these gaps in our scientific understanding of cone physiology, there are limitations in clinical DA measurement. Rod-mediated adaptation is a slow process, often requiring 30 to 40 minutes of testing time. This long duration causes fatigue, particularly in elderly populations [4, 13]. Moreover, psychophysical measurements are inherently subjective, while ERG typically requires contact electrodes, making it an invasive procedure.

Optoretinography (ORG) has emerged as a promising noninvasive imaging modality that provides an objective measure of cone or rod photoreceptor function with 3D cellular resolution. By employing an adaptive optics (AO) optical coherence tomography (OCT) system, ORG measures stimulus-evoked optical path length (OPL) changes in individual photoreceptor outer segments (OS) [28–32]. The elongation stage of the ORG response has been hypothetically attributed to phototransduction-related osmotic swelling of the outer segment [33, 34], hydration of the opsin proteins or the disc membrane they span [35], and/or mechanical force due to conformational changes in activated phototransductive enzymes [36].

In this study, we utilize high-resolution ORG to monitor photoreceptor dynamics *in vivo*, specifically focusing on the evolution of the ORG response during dark adaptation. The 3D cellular resolution of ORG allows us to isolate single photoreceptors and directly quantify cone-specific recovery processes. We present an efficient framework for DA measurement by ORG and propose two analytic methods to model ORG dynamics during the recovery phase, applying the model parameter time constant *τ* to represent the speed of recovery. This methodology provides new insight into the cellular mechanisms underlying dark adaptation and provides a practical method for evaluating functional deficits in retinal disease.

## 2 Results

### 2.1 AO-OCT visualizes single cone photoreceptors and reveals the phase of light reflected from their outer segments

In order to measure the cone photoreceptor’s light-evoked responses and their modulation by dark adaptation, the cones must be resolved in three dimensions. Figure 1a is a schematic representation of the 3D AO-OCT image, with a B-scan illustrated by the green box and areal views of the inner-outer segment junction (ISOS) and cone outer segment tips (COST) illustrated by the blue and red boxes, respectively. Magnified views of these projections are shown in Fig. 1b. The ISOS and COST projections reveal the quasi-periodic mosaic of cones, with one-to-one correspondence between the layers. Each pair of corresponding reflections represent light reflected from the ISOS and COST, which form the inner and outer boundaries of the cone outer segment (OS). Figure 1c depicts the OS, which consists of a stack of membraneous discs studded with the light-sensitive opsin proteins. Isomerization of the 11-cis-retinal chromophore within the opsin by an incident photon initiates phototransduction in the OS and the process of vision. The phase of the light waves reflected by the ISOS and COST, labeled *θ*_*ISOS*_ and *θ*_*COST*_, respectively, is used to infer the relative movement of these layers at the nanometer scale. This relative movement is the source of the optoretinographic (ORG) response, depicted in Fig. 1d. Details of the calculation and fitting of the ORG response are provided in the Methods.

**Fig. 1.**
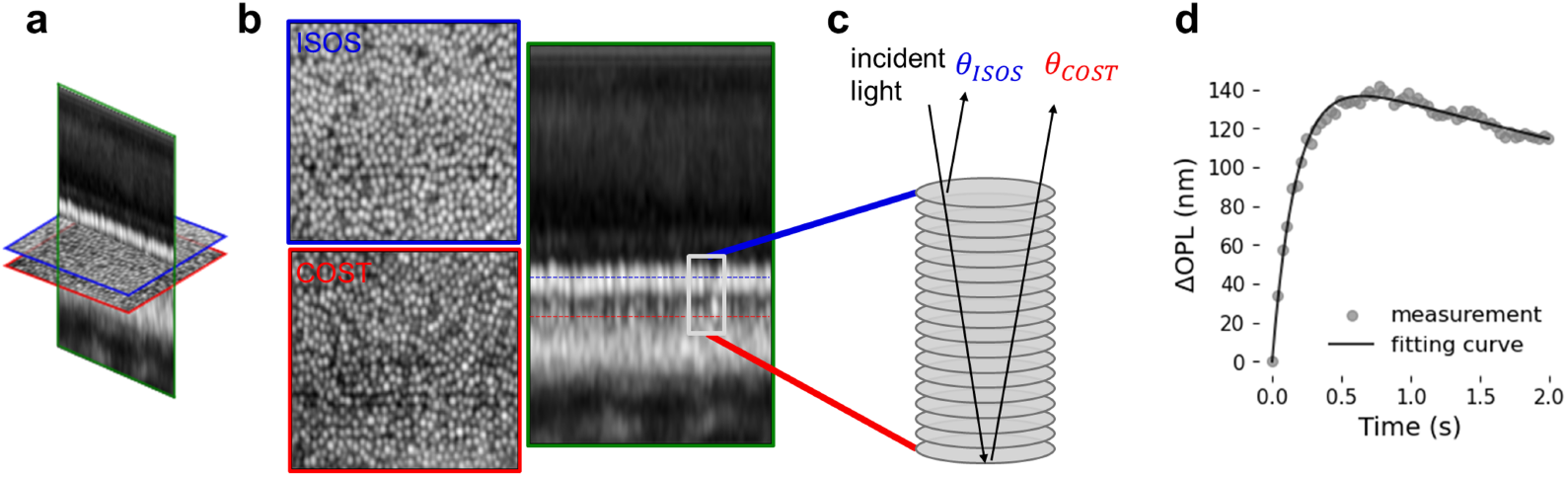
Cones are visualized in 3D in volumetric AO-OCT images of the retina. (a) is a schematic representation of the 3D AO-OCT image, with a longitudinal section (B-scan) illustrated with a green box and two transverse sections (C-scans) illustrated with blue and red boxes. (b) shows these sections in detail, revealing the quasi-periodic structure of the cone photoreceptors. The transverse sections are at the levels of the inner-outer segment junction (ISOS) and cone outer segment tips (COST) of the cone photoreceptors, which form the boundaries of the outer segment (OS) which is the site of phototransduction. The stack of membraneous discs constituting the OS is depcted in (c), with the phase of the light waves reflected by these surfaces labeled *θ*_*ISOS*_ and *θ*_*COST*_. By monitoring the difference between these phases, nano-scale changes in the length of the OS can be calculated. An example response to a visible stimulus is shown in (d), consisting of an elongation stage and a late contraction stage.

### 2.2 The cones’ ORG responses to light stimuli change during dark adaptation

Figure 2 shows ORG responses during a 304-second dark adaptation trial of subject 2, averaged among hundreds of cones in the stimulated area. The red traces depict the change in the cone outer segment optical path length (ΔOPL) resulting from equal, weak ‘probe flashes’ delivered at sequential acquisition times. ΔOPL was computed by monitoring the difference between the phase of light waves reflected by the two ends of the OS and converting these angular values into geometric lengths (see Eq. 2). The initial green trace at *t* = 0 corresponds to the ORG response elicited by an initial ‘bleaching flash’ that bleached 67 % of photopigment. The tone of each red trace indicates the increasing temporal delay between the initial 67 % bleach and each probing measurement.

**Fig. 2.**
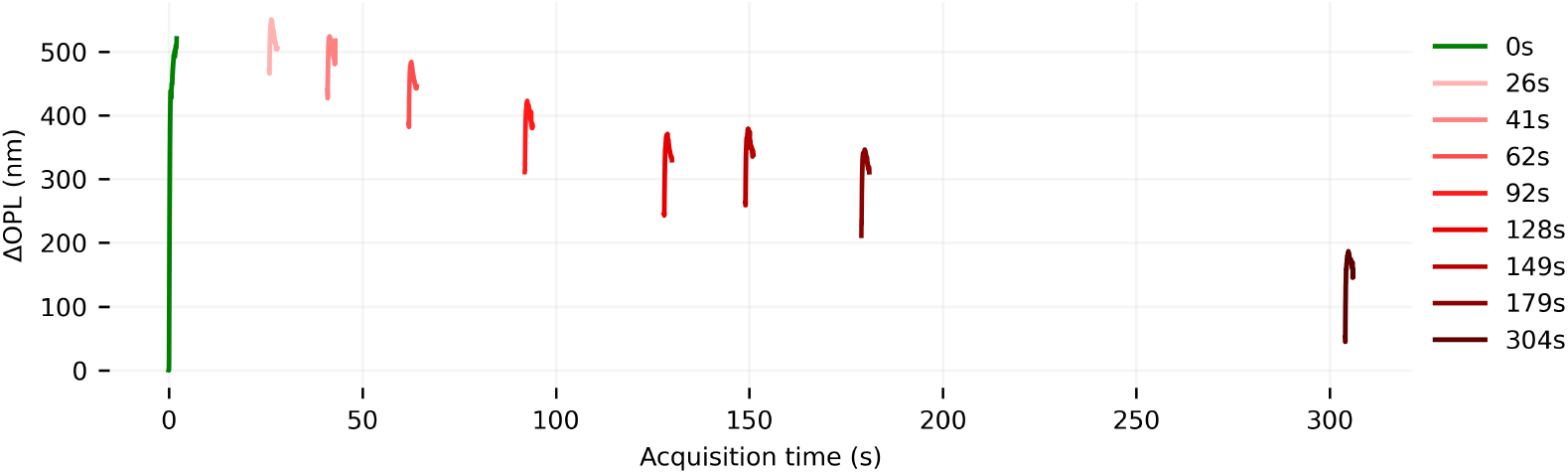
Responses of cone photoreceptors to light stimuli, recorded during a period of dark adaptation lasting 304 s. Light stimuli cause deformations of the cone photoreceptor outer segment (OS). These deformations include an elongation lasting hundreds of milliseconds, followed by a slower contraction. Plotted in green (*t* = 0) is the cones’ response to the initial bleaching flash, designed to isomerize 67 % of photopigment. Plotted in shades of red are responses to much dimmer probe flashes (8 %), at various times during the process of dark adaptation. During dark adaptation, two effects are observed: the baseline (pre-stimulus) OS length falls slowly over 304 s, while the amplitude of the responses (relative to baseline) rises. These responses were recorded during a single trial in subject 2. Fig. 3a shows responses like these, superimposed on a single 2 s time scale.

Figure 3 shows superimposed ORG traces from two of the subjects. The dotted red traces depict the change in the cone outer segment (OS) optical path length (ΔOPL) resulting from the probe flash stimulus delivered at *t* = 0. The tone of each red trace indicates the effective delay between its measurement and the initial bleaching flash. Because the cone responses were acquired in series after a single bleaching flash, rebleaching by probe flashes required correction of the time elapsed after the bleaching flash (described in Methods). Corrected (or effective) adaptation times are referred to as *t*_adapt_, and listed in the figure legend. From these traces it is evident that the ORG response of the cones becomes stronger as they adapt to the dark.

**Fig. 3.**
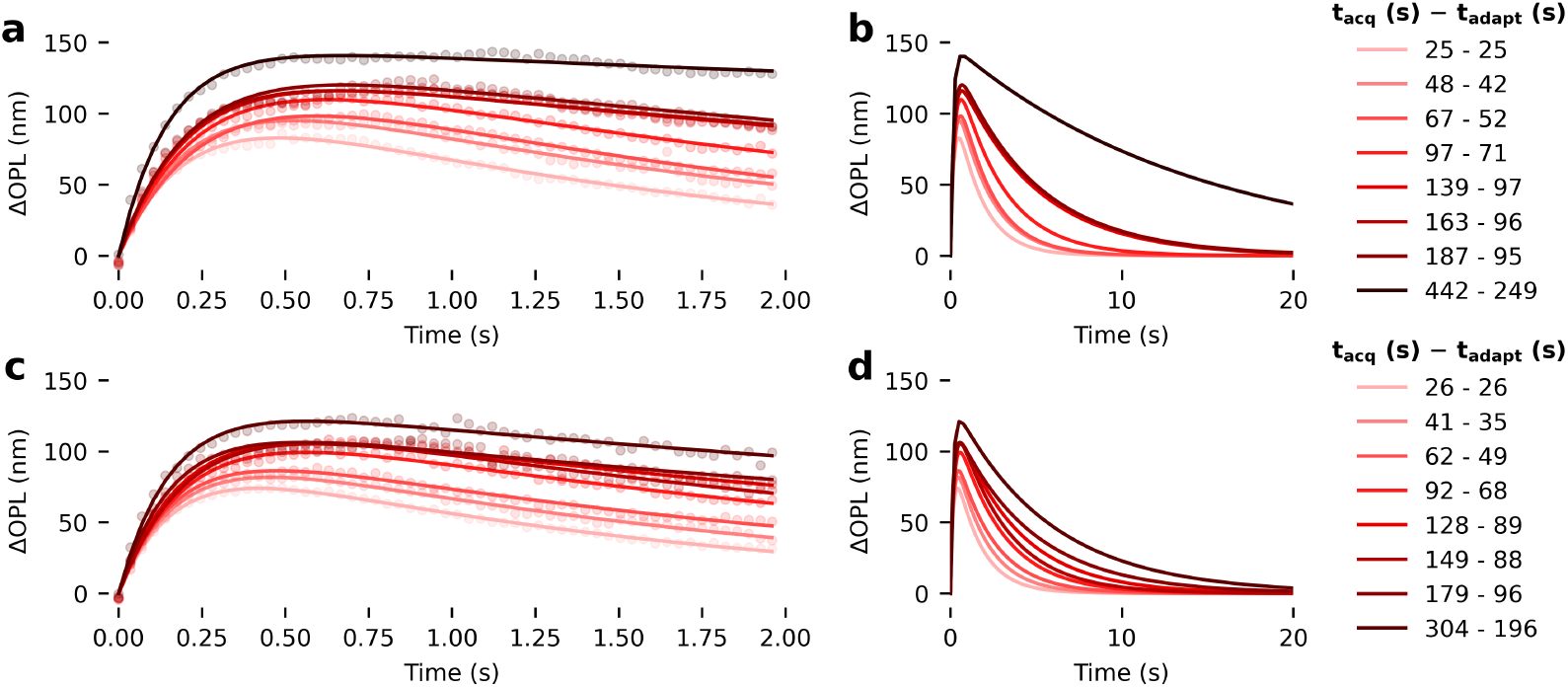
Cone photoreceptor responses to light stimuli become stronger during dark adaptation. The dotted lines shown in **a** and **c** are families of cone photoreceptor ΔOPL responses in two of the subjects, with varying amounts of adaptation time after a bright bleaching flash (*t*_adapt_, indicated in the legend). Because these responses were all recorded during a single period of dark adaptation, *t*_adapt_ was corrected for rebleaches, by subtracting the expected rebleach regeneration time from the real time. Broadly, the responses can be seen to become stronger as *t*_adapt_ increases. In particular, the maximum elongation ΔOPL_max_ and time-to-peak *t*_pk_ increase with *t*_adapt_, while the rate of late contraction *τ*_*b*_ decreases. Responses were fit with an overdamped oscillator model (Eq. 4), and the resulting fits to the data are plotted with solid lines. Fitting parameters permitted extrapolation of the responses, illustrated in panels **b** and **d**, which highlights the effect of dark adaptation on *τ*_*b*_.

### 2.3 ORG responses were parameterized in order to study their evolution

The ΔOPL plots in Fig. 3 exhibit two distinct stages of outer segment length change: (1) In the rising phase, the maximum OPL change (ΔOPL_max_) and the duration of outer segment elongation both increased with longer dark adaptation. For example, in subject 1, the maximal outer segment length change at 25 s of adaptation was approximately 80 nm, and it increased by ~75 % to ~140 nm at 442 s. (2) In the falling phase, the late contraction rate of the outer segment became noticeably slower with longer adaptation, as observed from the flattening ORG traces. The earliest reported stage of the cone ORG response is an initial rapid contraction, occurring within the first 10 ms [29, 37] and attributed to the early receptor potential (ERP) [34, 38], but the imaging rate employed here (28 Hz) was too low to detect it and the present work addresses only the elongation and late contraction stages described above.

Ultimately we were interested in quantifying the apparent changes in the cone ORG responses during the course of dark adaptation, which requires parameterization of the curves shown in Figs. 2 and 3. To do this, cone ΔOPL responses were fit with a model based on an overdamped resistor-inductor-capacitor (RLC) oscillator [39], shown in Eq. 4. These fits are depicted in Fig. 3 with solid lines. The resulting model parameters were then used to extrapolate ORG responses up to 20 s for these two subjects, as shown in Fig. 3b. To quantify the amplitude of OS elongation, we evaluated (1) the maximal OS elongation, ΔOPL_max_, directly from the measurements and (2) the peak derived from the RLC model fitted ORG curves, 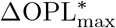. To quantify the temporal or dynamic characteristics of the responses, we evaluated two other parameters derived from the RLC model: (1) the late contraction time constant *τ*_*b*_, and (2) the time-to-peak *t*_pk_ denoting the duration of the initial outer segment elongation.

Preliminary inspection of the curves in Figs. 2 and 3 suggested that the ORG responses exhibit saturation, consistent with exponential decay. Generally, dark adaptation has been quantified through the time constant of exponential fits to the data, be they psychophysical [40] or electrophysiological [23]. For these reasons, we chose to perform exponential fits of the relationship between the model parameters ΔOPL_max_, 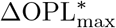, and *τ*_*b*_ as functions of adaptation time *t*_adapt_, which was calculated from the acquisition time *t*_acq_ as follows.

The experiment required acquisition of ORG measurements preceded by many, varying intervals of dark adaptation, amounting to hundreds of separate dark adaptation periods. To improve efficiency, we acquired multiple measurements during each period of dark adaptation, reducing the number of required trials by an order of magnitude. A necessary consequence of this design was that the ORG measurements, which required stimulus flashes, rebleached the photoreceptors and affected the resulting dynamics of adaptation. Thus the effective dark adaptation time, *t*_adapt_ in the preceding sections, was corrected for rebleaching. *t*_adapt_ was calculated by subtracting from acquisition time *t*_acq_ the estimated amount of time required to regenerate the photopigment bleached by the probe flashes, based on a standard exponential pigment regeneration model, shown in Fig. 7b. Details about this and alternative approaches are given in the Appendix.

### 2.4 The amplitude of OS elongation increases during dark adaptation

The two parameters related to the ORG amplitude, ΔOPL_max_ and 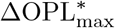, increased monotonically with adaptation time and exhibited saturation near the end of the trial, and were thus fit with an exponential function (Eq. 5), the speed of which is expressed by the resulting time constant *τ*.

As shown in Fig. 4, the data (Fig. 4a-c: ΔOPL_max_, Fig. 4d-f: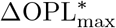) from 8-10 aggregated trials (colored circles, with color indicating trial number) in each of the three subjects were separately fitted with the exponential function in Eq. 5 (solid black lines). Model parameters *A* (scaling factor), *τ* (decay time constant), and *C* (asymptotic dark-adapted value) are listed on each plot. All fittings yielded a high coefficient of determination (*R*^2^ *>* 0.9).

**Fig. 4.**
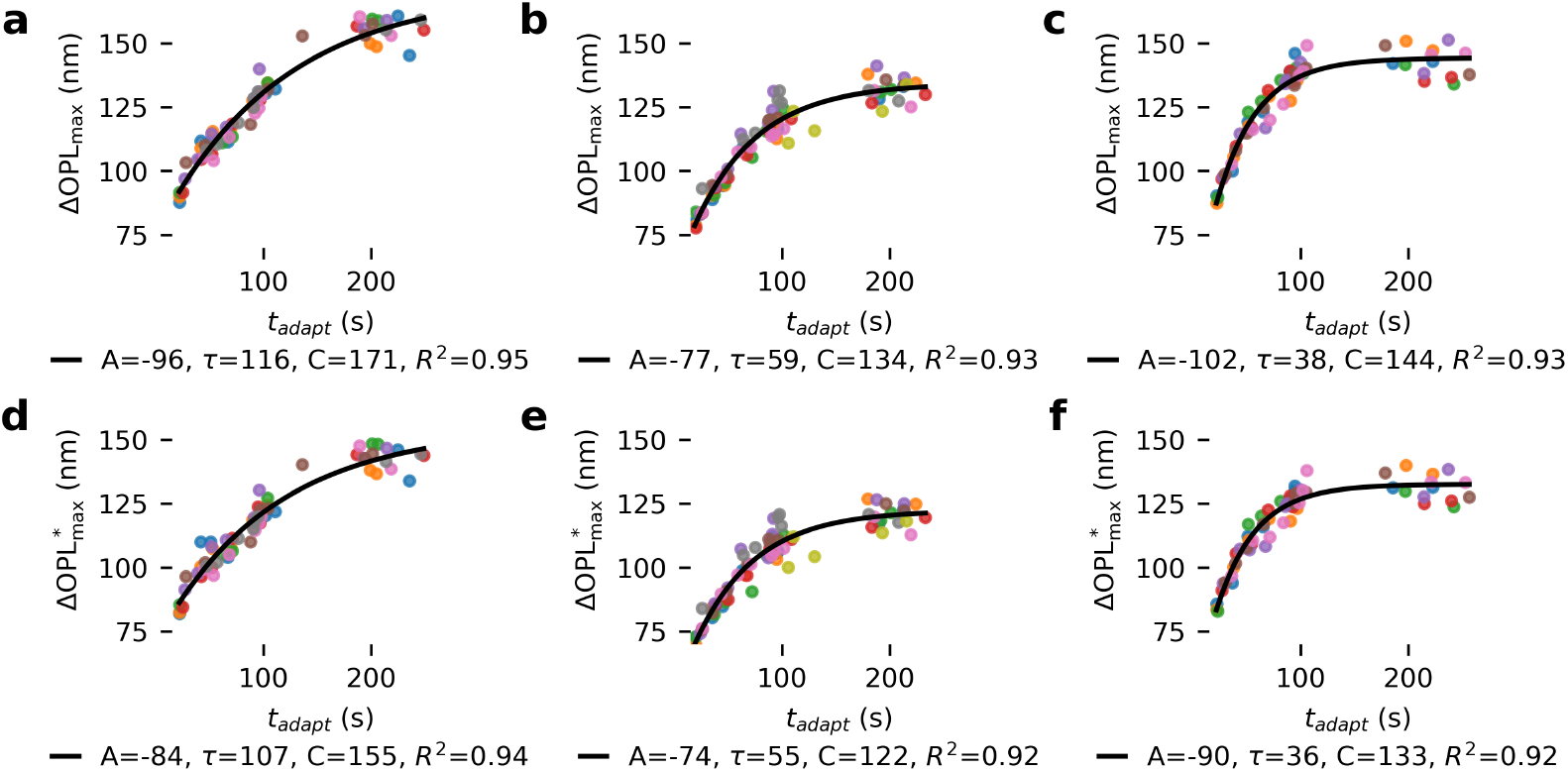
Cone ORG response amplitudes increase exponentially during dark adaptation. ORG response amplitudes are plotted as a function of dark adaptation time *t*_adapt_ for three subjects 1–3. **(a, b, c)** ΔOPLmax is the maximum cone OS elongation measured directly from ORG data. Responses were elicited by weak probing flashes (2.83 × 10^6^ photons*/*µm^2^). The response amplitudes increase and approach a saturation level as *t*_adapt_ increases. The corresponding maximum amplitudes derived from the model fit 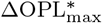 follow a similar decay trajectory, shown in panels **(d, e, f)**. Data from multiple trials are plotted and fitted together. The color of the markers indicates the trial. The overall decay trajectories were fit with an exponential model (Eq. 5), and the resulting fit is plotted with solid black lines. The parameters of each fit are printed below each panel. *A* indicates the decay amplitude, *τ* is the decay time constant in seconds, *C* denotes the asymptotic saturation level, and *R*^2^ indicates the coefficient of determination.

Given the similarities of ΔOPL_max_ and 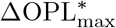, we expected similar fitting results for those two ORG amplitude metrics, and that was observed in the data shown in Fig. 4. The decay time constants *τ* for the two metrics were closely matched:116 s for ΔOPL_max_ vs. 108 s for 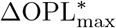 in subject 1, 59 s vs. 55 s in subject 2, and 38 s vs. 36 s in subject 3. The saturation level *C* in 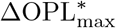 was consistently lower than in ΔOPL_max_, because ΔOPL_max_ represents an average of five maxima, which increases its overall value.

Interestingly, the time constants *τ* and peak amplitudes ΔOPL_max_ varied considerably across subjects: 116 s and 171 nm for subject 1, 59 s and 134 nm for subject 2, and 38 s and 144 nm for subject 3. The time constant differed by nearly a factor of three between subjects 1 and 3.

### 2.5 The time-to-peak of the cone response increases during dark adaptation while the late contraction rate decreases during dark adaptation

Fig. 5 shows the evolution of *t*_pk_ and *τ*_*b*_ as a function of adaptation time, *t*_adapt_, with exponential fits (Eq. 5) for subjects 1–3. The metric *t*_pk_ increased with *t*_adapt_, while *τ*_*b*_ decreased, which aligned with the observations in the representative ORG curves shown in Fig. 3. The outer segment elongation duration *t*_pk_ exhibited an exponential increase (Fig. 5(a - c)), and upon fitting yielded time constants of 79 s, 45 s and 25 s for subjects 1-3, respectively, with coefficients of determination *R*^2^ ranging between 0.79 and 0.84. The exponential model also described the dynamic decrease in *τ*_*b*_ (Fig. 5(d - f)). The corresponding time constants were 66 s, 51 s, and 32 s, with *R*^2^ *>* 0.9.

**Fig. 5.**
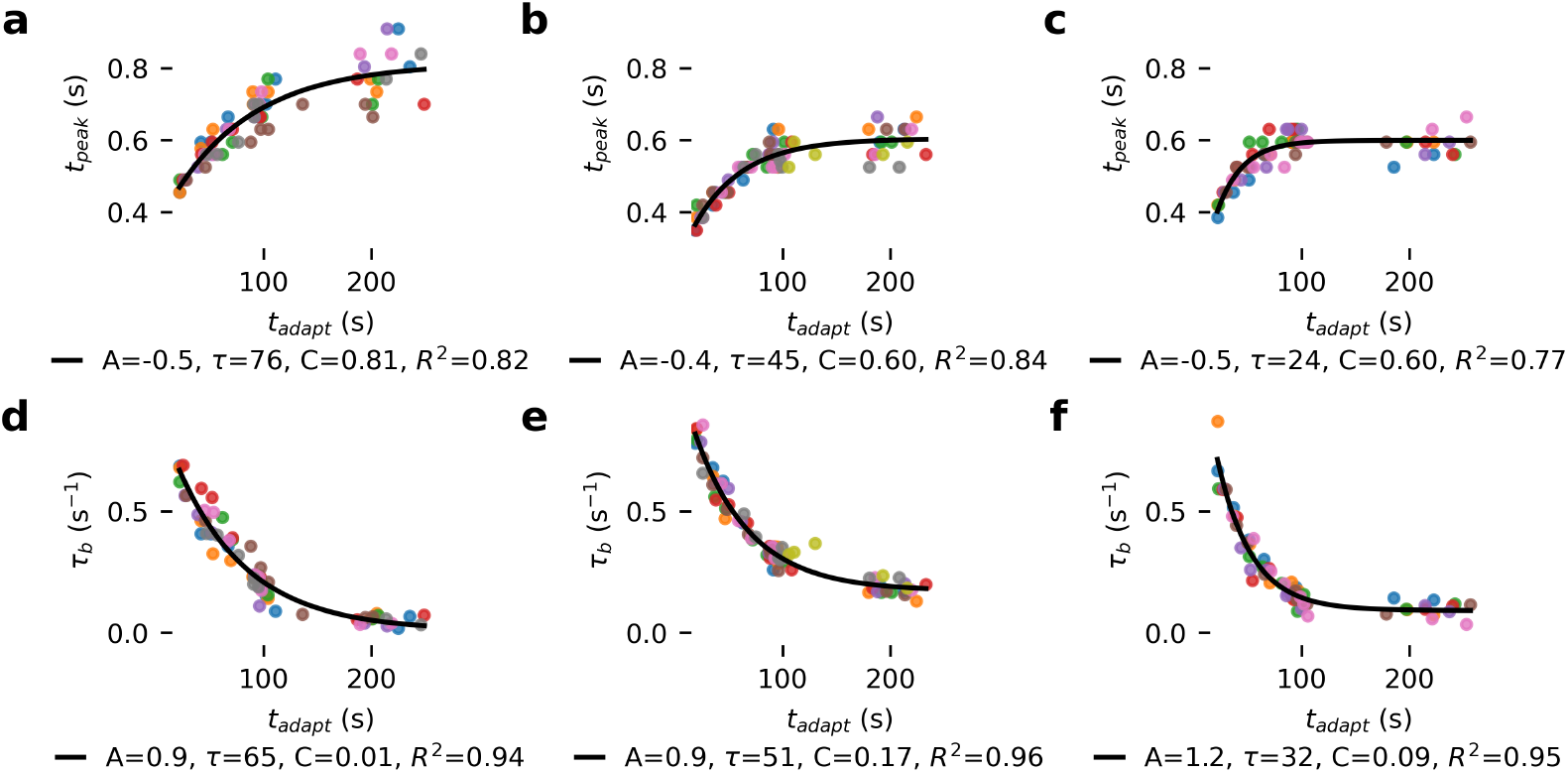
Cone ORG temporal dynamics recover exponentially during dark adaptation. **(a, b, c)** denote the time-to-peak (*t*_pk_) and **(d, e, f)** show the late contraction rate (*τ*_*b*_) from three subjects 1–3 from left to right, as functions of dark adaptation time (*t*_adapt_). Data points are color-coded to indicate 8-10 repeated trials. The overall decay trajectories were fit with an exponential model (Eq. 5), shown as solid black lines, with the corresponding fitting parameters displayed at the bottom of each panel.

### 2.6 ORG amplitude exhibited slower decay than ORG dynamics during dark adaptation

Table 1 summarizes the adaptation time constants obtained from exponential fitting (Eq. 5) for the ORG amplitude and dynamic parameters in three subjects.

**Table 1.** Summary of the decay time constants for cone ORG parameters during dark adaptation. The decay time constant *τ* (s) was derived through exponential fitting (Eq. 5) of cone response amplitudes (ΔOPL_max_ and 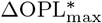) and temporal parameters (*t*_*pk*_ and *τ*_*b*_) as functions of dark adaptation time *t*_adapt_.

| Subject | $\Delta OPL_{max}$ | $\Delta OPL_{max}^*$ | $t_{pk}$ | $\tau_b$ |
| --- | --- | --- | --- | --- |
| 1 | 116 | 107 | 79 | 66 |
| 2 | 59 | 55 | 45 | 51 |
| 3 | 38 | 36 | 25 | 32 |
| Mean | 71 | 66 | 50 | 50 |

The ORG amplitude exhibited the slowest decay, with average adaptation time constants of 71 s and 66 s for the decay to baseline of ΔOPL_max_ and 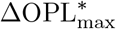, respectively. The decay of parameters related to dynamics, *t*_*pk*_ and *τ*_*b*_, was faster, with average adaptation time constants of 48 s and 49 s, respectively.

Across all ORG metrics, subjects 1 and 3 exhibited the slowest and fastest recovery, respectively. The differences between subject 1 and subject 3 were approximately two-to threefold across the four ORG metrics.

## 3 Discussion

This study establishes an objective, noninvasive method for quantifying human cone dark adaptation *in vivo* using optoretinography with adaptive optics OCT. By monitoring stimulus-evoked optical path length changes between the ISOS and COST layers at the single-cone level and parameterizing these responses, we quantified changes in both the amplitude and temporal dynamics of cone responses over a 400 s recovery period following a strong (67 %) photopigment bleach.

All ORG parameters, including response amplitude (ΔOPL_max_ and 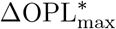), OS elongation duration (*t*_pk_), and late contraction rate (*τ*_*b*_), exhibited systematic, monotonic decay toward their baseline values during dark adaptation. To characterize these dynamics, we evaluated three analytic models: (1) an exponential model based on first-order pigment regeneration dynamics; (2) a rate-limited model of pigment pigment regeneration; and (3) a perturbed exponential model accounting for perturbations from repeated test flashes. Results from the first approach are presented in the Results above. Results from the other two methods are described in the Appendices.

Importantly, all three models yielded consistent recovery trends across subjects and ORG parameters. The fitted decay constants (*τ* or *v*) demonstrates two robust characteristics of cone-mediated dark adaptation: (i) the amplitude metric ΔOPL_max_ recovered more slowly than the temporal metrics *t*_pk_ and *τ*_*b*_, which may indicate that partly distinct mechanisms underlie the two; and (ii) substantial inter-subject variability (approximately two-to threefold differences), indicating physiological diversity in cone pigment regeneration or phototransduction recovery kinetics.

These findings support the interpretation that ORG dynamics reflect not only photopigment regeneration in canonical and cone-specific pathways [2, 5, 13], but potentially also downstream processes in the phototransduction cascade. Specifically, discrepancies between amplitude- and time-based metrics may arise from alterations in protein deactivation and recycling rates. Elucidating these components, such as the roles of specific proteins and the kinetic properties of visual cycle enzymes, will necessitate targeted experiments under controlled manipulations, potentially utilizing murine genetic models [13, 41–43]. Practically, the strong agreement among the three modeling approaches confirms that ORG provides a reliable, quantitative measure of dark adaptation. The approach is objective, rapid (~7 min per trial), noninvasive, and resolved at the level of individual photoreceptors.

Additionally, we used the AdaptDx dark adaptometer (OpZira, Inc.) to evaluate the psychophysical visual sensitivity recovery in the same three subjects. To quantify the rate of cone dark adaptation, an exponential model was fit to the sensitivity measurements obtained during the initial cone-mediated recovery phase prior to the rod intercept (up to 250 s), as shown in Fig. 6. The psychophysical cone recovery time constants *τ* for subjects 1–3 were 46 s, 38 s, and 31 s, respectively. Interestingly, these perceptual recovery time constants were shorter than the corresponding ORG parameter recovery time constants measured by Method 1 (Table. 1) and Method 3 (Table. C1). This discrepancy suggests that post-receptoral amplification or higher-order cortical processing may accelerate perceptual visual recovery beyond the biochemical processes occurring within the retina.

**Fig. 6.**
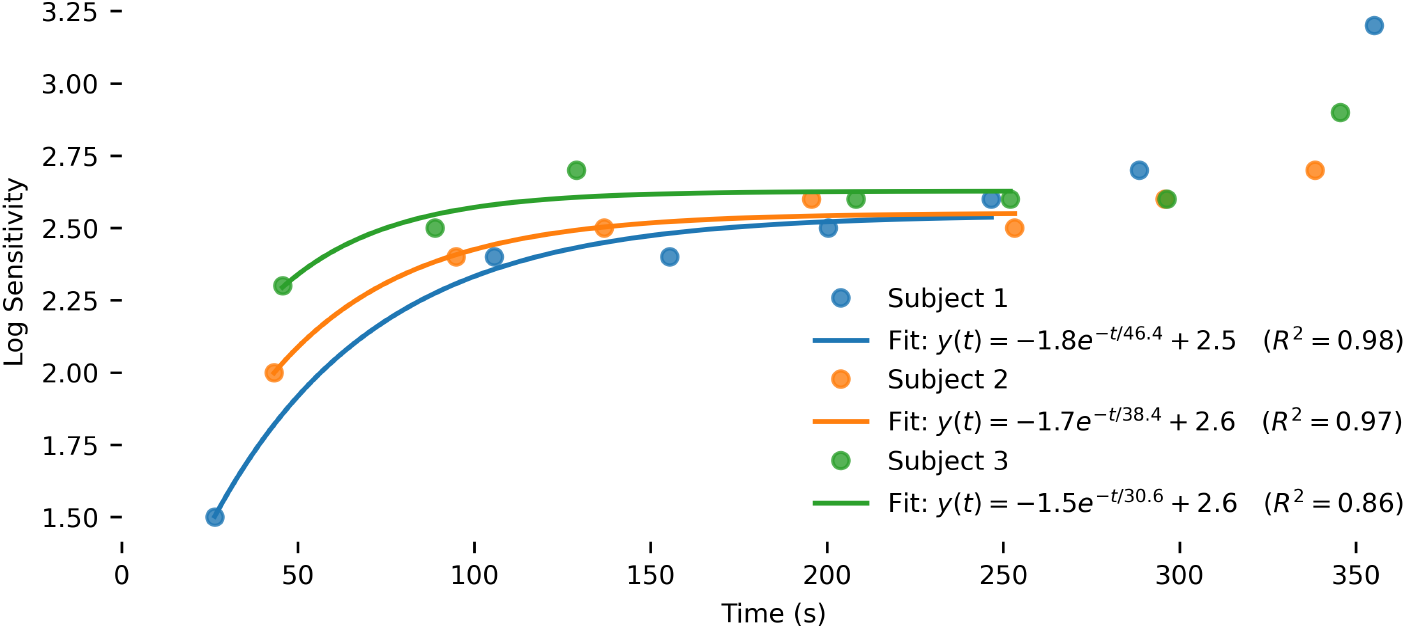
Psychophysical visual sensitivity recovers exponentially during dark adaptation. Log sensitivity measured by the AdaptDx dark adaptometer (OpZira, Inc.) is plotted as a function of dark adaptation time for three subjects 1–3. The early recovery trajectories for the cones were fit with an exponential model (*y*(*t*) = *Ae*^−*t/τ*^ + *C*), shown as solid colored lines, with the corresponding fitting equations and coefficients of determination (*R*^2^) displayed in the legend.

**Fig. 7.**
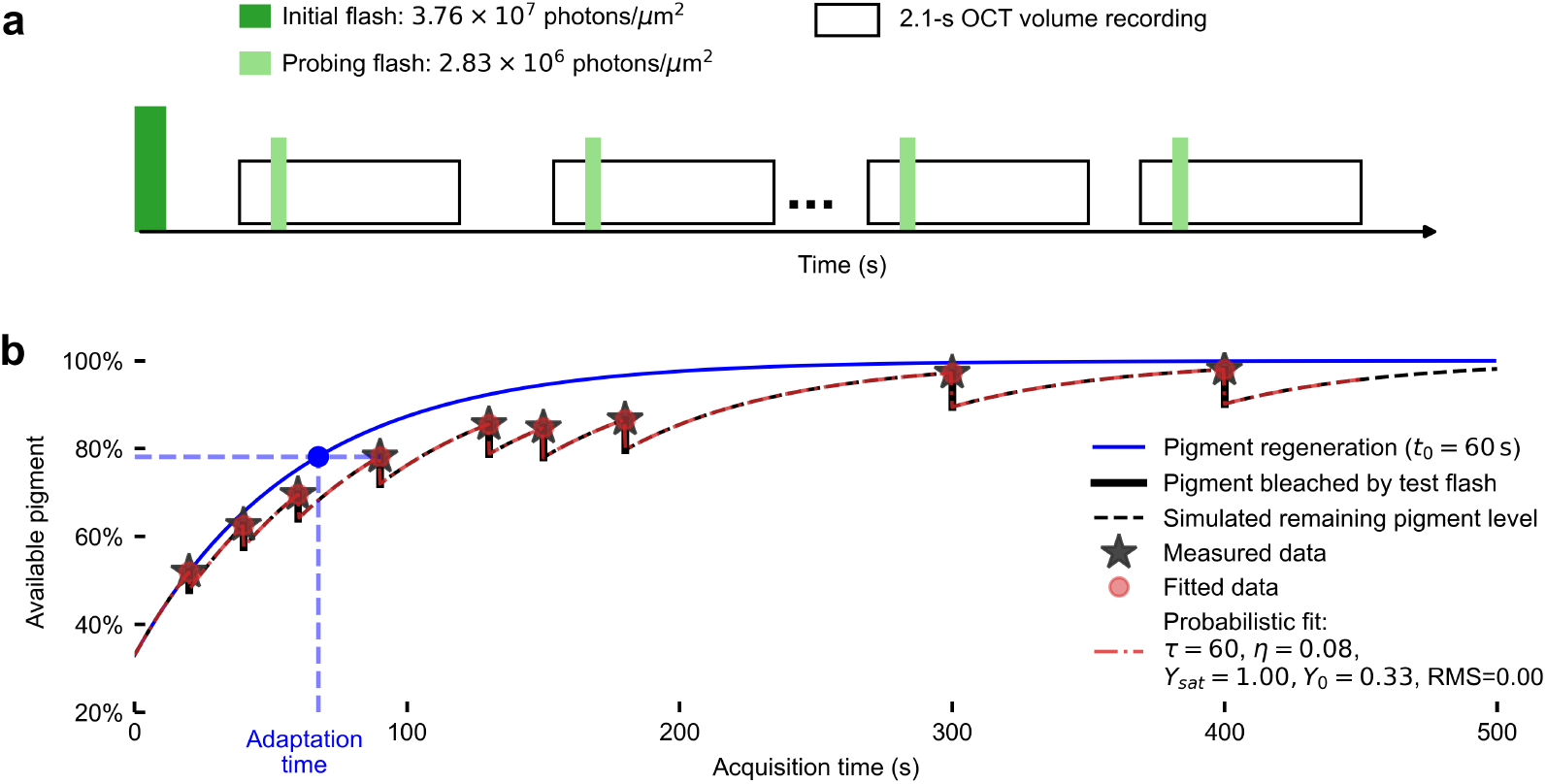
Experimental protocol of measuring dark adaptation with ORG. **a** Schematic of OCT image acquisition and flash delivery. **b** Modeled exponential pigment regeneration [17] following a 67 % bleach (Eq. A1 with *τ* = 60 s; solid blue curve). The dashed black curve depicts the predicted regeneration trajectory when perturbed by repeated ORG probing flashes (vertical black lines). Black stars mark the ORG recordings at nine predefined acquisition time points (*t* = [20, …, 400] s) at which test flashes (2.83 × 10^6^ photons*/*µm^2^) were delivered. To derive the effective adaptation time (blue marker), the photopigment fraction at each measurement point is mapped onto the unperturbed regeneration curve. Red dots and red dashed curves represent the perturbation model fit to the simulated regeneration data, accounting for rebleaching by the probe flashes.

Future advancements in motion correction, single-photoreceptor tracking stability [44], and protocol optimization will further enhance robustness, expand the framework to include both cone- and rod-mediated dark adaptation [41, 45, 46], shorten imaging sessions, and facilitate translation to clinical populations. Given that RPE function, retinoid transport, and phototransduction defects can alter dark adaptation prior to visible structural changes [4, 13, 16], ORG holds promise as a highly sensitive biomarker for early dysfunction in retinal pathologies, including AMD, inherited retinal degenerations, and vitamin A deficiency, and may enable longitudinal monitoring of emerging therapies targeting the visual cycle.

In summary, dark-adaptation ORG offers a powerful new window into mechanisms of dark adaptation, bridging microscopic photoreceptor dynamics with clinically relevant functional assessment. This work establishes a foundation for future investigation and diagnosis at the earliest stages of retinal dysfunction related to dark adaptation.

## 4 Methods

### 4.1 Imaging system

The AO-SS-OCT system used in this study is described in detail in our previous work [47]. It comprised an OCT system with a Fourier domain mode-locked (FDML) laser (FDM-1060-750-4B-APC, OptoRes GmbH, Munich, Germany) operating at an A-scan rate of 1.64 MHz. The AO subsystem incorporated a pupil-conjugated deformable mirror (DM-97-15, ALPAO) and a custom-made Shack–Hartmann wavefront sensor [48]. The FDML laser power at the cornea was 1.8 mW. A superluminescent diode at 840 nm (Superlum Diodes Ltd, Cork, Ireland) served as the wavefront beacon, with power at the cornea of 150 µW. The system measured and corrected aberrations over a 6.75 mm pupil at a closed-loop rate of 10 Hz using custom AO control software [48], providing a theoretical lateral resolution of 3.2 µm. A resonant scanner (SC-30, Electro-Optical Products Corp., Ridgewood, New York) oscillating at 5 kHz provided horizontal scanning, while a slower galvanometer scanner controlled vertical scanning. This raster scanning configuration, together with the FDML A-scan rate of 1.6 MHz, enabled a 28 Hz OCT volume rate over 1° × 1° field of view (FOV). Each OCT volume consisted of 160 A-scans per B-scan and 172 B-scans, corresponding to sampling intervals of 1.875 µm and 1.744 µm in the fast and slow scanning dimensions. These fall short of the minimum sampling frequency of 1.6 µm suggested by the lateral resolution of 3.2 µm, a compromise we accepted in order to maintain a 1° FOV.

A fiber-coupled LED (M565F3, Thorlabs, New Jersey) centered at 555 nm with 20 nm bandwidth served as the stimulus flash, producing light that bleaches L and M cones equally [49, 50]. A 4-f optical relay conjugated the retina with an iris in the stimulus channel, allowing adjustment of the stimulus area. The stimulus and fixation channels were combined using a dichroic mirror. A custom Badal system [51] was incorporated into both channels to provide defocus compensation.

The computation of the photopigment bleach level is detailed in [52, 53]. The stimulus area was 1.53° in diameter to fully cover the 1°× 1° imaging FOV. Two bleach levels were used in this study: a bleaching flash condition (67 %) with a photon density of 3.76 × 10^7^ photons*/*µm^2^, and a probe flash condition (8 %) with a photon density of 2.83 × 10^6^ photons*/*µm^2^.

### 4.2 Image acquisition

Three male subjects (ages 35-51 years), with no known retinal diseases, were dilated and cyclopleged using topical phenylephrine (2.5 %) and tropicamide (1.0 %). A bite bar and forehead rest were used to stabilize the eye pupil during imaging. A calibrated LCD screen was used to display the fixation target. Before imaging, subjects underwent 5 min of dark adaptation by staying in a dark room with the eye to be imaged covered by an eye patch.

All experimental procedures complied with the Declaration of Helsinki and were approved by the Institutional Review Board of the University of California, Davis. Written informed consent was obtained from each participant after the study procedures and potential risks were fully explained. The combined illumination from the three sources met ANSI laser safety standard [54].

Images were acquired at 3° nasal 4° superior (3N4S) following the protocol illustrated in Fig. 7**a**. Each trial was initiated by delivering a 35 ms bright flash calibrated to bleach 67 % of L- and M-photopigments. Over the following 400 s, ORG recordings were collected at nine known time points. Each recording was acquired at 28 Hz volume rate over 2.1 s, consisted of 60 volumes, with a 10 ms probing flash (2.83 × 10^6^ photons*/*µm^2^) delivered after the second volume. For each subject, 8-10 repeated trials were conducted.

### 4.3 Data processing

The ORG processing pipeline has been detailed in our previous work [55]. Following OCT volume registration [28], the phase difference between ISOS and COST was extracted for individual cones at each acquisition time *t*_*n*_ and converted into optical path length change ΔOPL in the cone OS [30]:

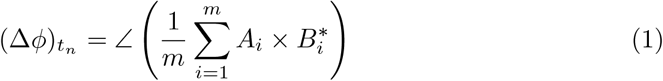

where *A* is the complex OCT signal measured at COST layer, *B*^∗^ is the complex conjugate of the OCT signal measured at ISOS, and *m* is the number of A-scans recorded within each single cone. In the present study, we segmented cones using 7 × 7 pixel region. From this region containing the cone, the 9 brightest A-scans (m=9) were selected at each time point for ORG processing. The change in optical path length (ΔOPL) in cone OS was calculated based on the imaging wavelength *λ* using

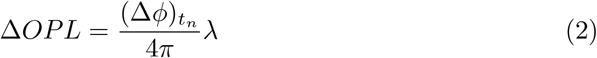

To quantify the overall cone outer segment elongation, an overdamped RLC model [39] was fit to the measured ΔOPL as below:

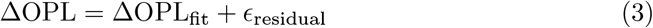

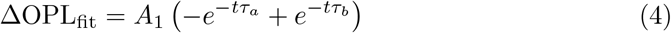

where 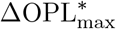 is the model-fitted optical path length change, and *ϵ*_residual_ is a residual term that accounts for measurement noise and unmodeled physiological effects.

Four ORG-derived parameters were quantified from each fit to evaluate the overall ORG response: (1) ΔOPL_max_ (unit: nm): the maximum optical path length change in cone OS, calculated as the average of the five highest values of ΔOPL; (2) 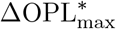 (unit: nm): the peak amplitude derived from the RLC model fit (Eq. 4) ΔOPL_fit_; (3) *τ*_*b*_ (unit: s^−1^): the parameter from the fitting model (Eq. 4) describing the late, slow contraction rate of the outer segment [39]; (4) *t*_pk_ (unit: second): time to peak by solving t for the first zero of the model’s derivative *d*(ΔOPL_fit_)*/dt* = 0. Each parameter was averaged across all 200 − 800 segmented cones at each acquisition time within each trial.

### 4.4 Analytic models for data analysis

Three methods were employed to characterize the ORG dynamics during dark adaptation.

The first is based on an exponential photopigment regeneration model with a time constant of 60 s. For each ORG measurement, the available pigment level was mapped onto the unperturbed pigment regeneration curve to determine the effective dark adaptation time, *t*_adapt_, as demonstrated in the Fig. 7b. Full mathematical details are provided in Appendix A. The ORG parameters were then fitted as a function of *t*_adapt_ using an exponential form:

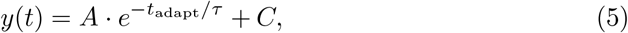

where *τ* represents the decay time constant and *C* denotes the asymptotic saturation level.

The second method is analogous to the first, but using a rate-limited model of pigment regeneration [25, 26]. Unlike simple first-order exponential fitting, this model assumes that the recovery is limited by retinoid delivery to the outer segment and has provided improved fits for cone-driven ERG a-wave recovery [25]. To test if this rate-limited framework better described ORG dynamics in dark adaptation, we applied this rate-limited model to simulate the pigment regeneration and fit into the ORG parameters trajectories during dark adaptation, as detailed in Appendix B. It is not depicted in Fig. 7.

The third method fits the data to a perturbed exponential model, without assumptions about the pigment regeneration time constant. This approach evaluates the ORG parameters directly as a function of the experimental acquisition time (*t*_acq_). Conceptually, the model accounts for two parallel processes: the continuous exponential recovery of the photoreceptor in the dark, and the discrete perturbations induced by each test flash.

Every time a probing flash is delivered, the model applies a proportional drop to the recovery process, representing the transient bleaching effect. Immediately following this drop, the dark-adaptation recovery resumes toward its final dark-adapted state. Iterating this drop-and-recover sequence across all acquisition times generates a “stepped” decay trajectory. The model parameters were optimized by minimizing the root-mean-square (RMS) error between this predicted trajectory and the measured ORG metrics. This fit yields four interpretable parameters: (1) *τ* (s), the intrinsic decay time constant; (2) *η*, the magnitude of the flash-induced perturbation; (3) *Y*_sat_, the asymptotic dark-adapted saturation level; and (4) *Y*_0_, the initial post-bleach value at 0 s. The full mathematical formulation of this iterative process is detailed in Appendix C.

To validate this method, the model was fit to the simulated pigment dynamics shown in Fig. 7. Using the discrete pigment levels at the predefined time points (black stars) as input measurements, the model predictions (red dots) accurately reproduced the underlying theoretical trajectory (red dashed line vs. black dashed line). The fit yielded parameters consistent with the simulation inputs: *τ* = 60 s for regeneration, *η* = 0.08 (8 %) for flash-induced perturbation, *Y*_sat_ = 1, and *Y*_0_ = 0.33 (reflecting the initial 67 % bleach).

## Data availability

The processed datasets generated and analyzed during the current study, including the optical path length (ΔOPL) responses, RLC model fitting ORG parameters 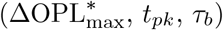 are publicly available in the Zenodo repository [56]. Raw volumetric AO-OCT imaging datasets are available from the corresponding author upon reasonable request.

## Code availability

The custom analytical code used to analyze the ORG parameters, including the three analytic methods described in Section 4.4, is available in the Zenodo repository [56]. The scripts are written in Python 3 and rely on standard open-source libraries.

## Author contributions

Study conception and design: Y.C., R.J.Z., and R.S.J.; Imaging system preparation: Y.C. and M.M.B.; Data collection: Y.C.; Data analysis and pipeline development: Y.C., R.S.J., R.J.Z., A.Z. and R.M.; Data interpretation: Y.C., R.J.Z., and R.S.J.; Drafting of the manuscript: Y.C. and R.S.J. All authors contributed to editing the manuscript.

## Acknowledgements

This work was supported by NIH grants R01-EY-034340, R01-EY-033532, R01-EY-031098, R01-EY-26556, and P30-EY-012576. The authors would like to acknowledge: the advice of colleagues including John S. Werner, Omar Mahroo, and Hannah Smithson; management of the IRB and clinical coordination by Susan Garcia; and the support, financial and otherwise, from the UC Davis Department of Ophthalmology and Vision Science.

## Ethics declarations

The authors declare no competing interests.

## Appendix A Method 1: Calculating effective adaptation time *t*_adapt_ using an exponential model of pigment regeneration

The first approach maps acquisition time (*t*_adapt_) onto a theoretical pigment regeneration model to derive an ‘effective adaptation time’ (*t*_adapt_), against which ORG parameters are fitted using an exponential function (Eq. 5), as shown in Fig. A1.

Because our experimental protocol utilizes sequential probing flashes to measure the optoretinogram during a single period of dark adaptation for around 5 min, there are hypothetically two processes happening at the same time: (1) a small fraction of photopigment bleach by the probing flashes; (2) photopigment regeneration in dark. Consequently, the actual available pigment at the time of data acquisition (*t*_acq_) is slightly lower than it would be in a perfectly unperturbed, dark-adapted eye. To account for this, we computed an effective adaptation time (*t*_adapt_), defined as the time it would take an unperturbed eye to reach the exact pigment level present at *t*_acq_.

The regeneration of human cone pigments in darkness was modeled as an exponential decay process of the bleached fraction [17]:

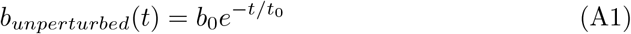

where *b*_*unperturbed*_(*t*) is the theoretical fraction of bleached pigment at time *t* in an unperturbed dark state, *b*_0_ is the initial high bleach level (67 %), and *t*_0_ is the pigment regeneration time constant (assumed to be 60 sec [17]), defined as the time required for the bleach level to decay to 1*/e* of its initial value. The corresponding unbleached pigment is therefore *p*_*unperturbed*_(*t*) = 1− *b*_*unperturbed*_(*t*), as shown by the blue curve in Fig. A1.

By incorporating the cumulative effects of continuous pigment regeneration and discrete flash-induced effective pigment depletion, we iteratively simulated the available pigment across the sequence of probing flashes as indicated by the dashed black line in Fig. A1. The effective pigment depletion produced by a probing flash at any given time, *b*_*eff*_ (*t*), is assumed to be proportional to the available photopigment (*p*(*t*)) at that exact moment:

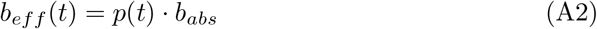

where *b*_*abs*_ is the bleach fraction produced by the test flash in a fully dark-adapted condition (8 % for a photon density of 2.83 × 10^6^ photons*/*µm^2^). Each probing flash transiently drops the available pigment level, after which exponential regeneration resumes until the next flash.

The remaining pigment fraction *p*(*t*_*acq*_) at each acquisition time *t*_acq_ is then mapped onto the ideal, unperturbed regeneration curve to determine *t*_adapt_:

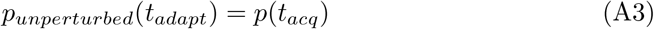

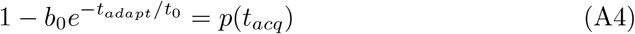

Solving for *t*_*adapt*_ yields the direct conversion formula used in this study:

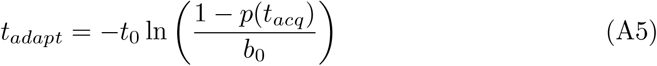

### A.1 Effect of pigment regeneration time constant

In Method 1, the effective adaptation time *t*_adapt_ was derived by mapping each ORG measurement onto an exponential cone pigment regeneration curve (Eq. A1). In the main analysis, we set the pigment regeneration time constant to *t*_0_ = 60 s [17, 18]. Because this choice directly affects the mapping from acquisition time to *t*_adapt_, we evaluated how sensitive the fitted ORG adaptation time constants *τ* are to the assumed value of *t*_0_.

To this end, we recomputed *t*_adapt_ for a family of pigment regeneration curves with *t*_0_ ranging from 30 to 200 s and refit all ORG metrics using the same exponential model (Eq. 5). For each subject and each metric, this yielded a set of fitted adaptation time constants *τ* (*t*_0_). Figure A2 summarized the dependence of *τ* on the pigment regeneration time constant *t*_0_ for four ORG metrics and three subjects.

**Fig. A1.**
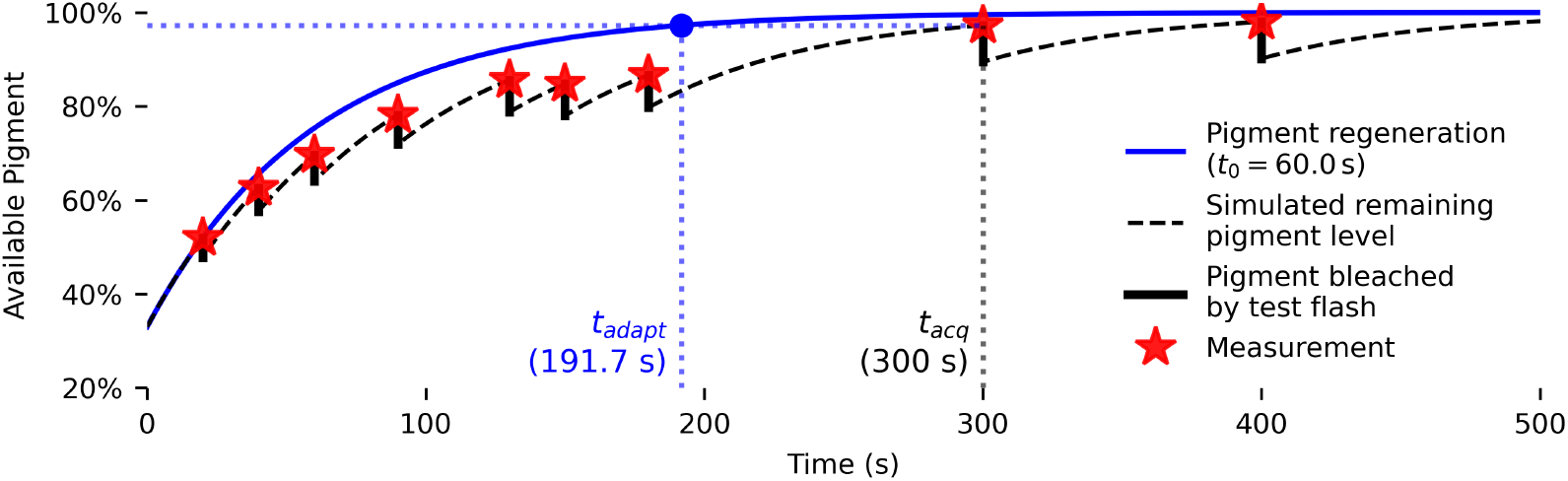
Calculation of the effective adaptation time (*t*_*adapt*_). The solid blue curve illustrates the ideal, unperturbed exponential pigment regeneration (*t*_0_ = 60 s) following an initial 67 % bleach (*b*_0_ = 0.67) at *t* = 0. The dashed black line represents the simulated trajectory of the available pigment level, accounting for the sequential effective pigment depletion fraction caused by 2.83 × 10^6^ photons*/*µm^2^ probing flashes delivered at *t* = [20, 40, 60, 90, 130, 150, 180, 300, 400] seconds, which is marked by the vertical black lines. To calculate *t*_adapt_ for any given ORG measurement (e.g., at acquisition time *tacq* = 300*s*), the available pigment fraction (red star) is projected horizontally onto the ideal regeneration curve (blue dot). The corresponding time on the x-axis represents *t*_adapt_ = 191.7*s*, effectively correcting the recovery timeline for the cumulative pigment depletion induced by previous probing flashes.

Across the full range of *t*_0_ values, the variation in *τ* was modest for all subjects and metrics. For example, for ΔOPL_max_ in subject 1, the fitted time constant decreased from *τ*_max_ = 116 s at *t*_0_ = 60 s to *τ*_min_ = 93 s at *t*_0_ = 200 s, corresponding to a change of approximately 20%. Similar behavior was observed for the other ORG metrics and subjects: the relative variation of *τ* as a function of *t*_0_ remained within ~ 20% in all cases. Importantly, the rank ordering of subjects (subject 1 consistently slowest, subject 3 consistently fastest) and the relative differences between amplitude and temporal metrics were preserved over the entire tested range of *t*_0_.

Thus, while the absolute numerical values of *τ* depend to some extent on the assumed pigment regeneration constant, the inter-subject differences and the qualitative relationships among ORG metrics are robust to this choice. The observed sensitivity of *τ* to *t*_0_ is clearly smaller than the inter-subject variability, suggesting that our main findings are not critically dependent on the exact value assumed for the cone pigment regeneration time constant.

## Appendix B Method 2: Calculating effective adaptation time *t*_adapt_ using a rate-limited model of pigment regeneration

Similarly, we evaluated a rate-limited model of cone pigment recovery proposed by Mahroo *et al*. [25, 26]. In this model, the overall recovery is limited by the delivery of 11-*cis*-retinoid to the photoreceptor outer segments rather than by simple first-order photopigment kinetics, providing improved fits to cone-driven ERG a-wave recovery over a range of bleach levels[25], motivating us to test whether it also better describes the recovery of ORG metrics measured here.

**Fig. A2.**
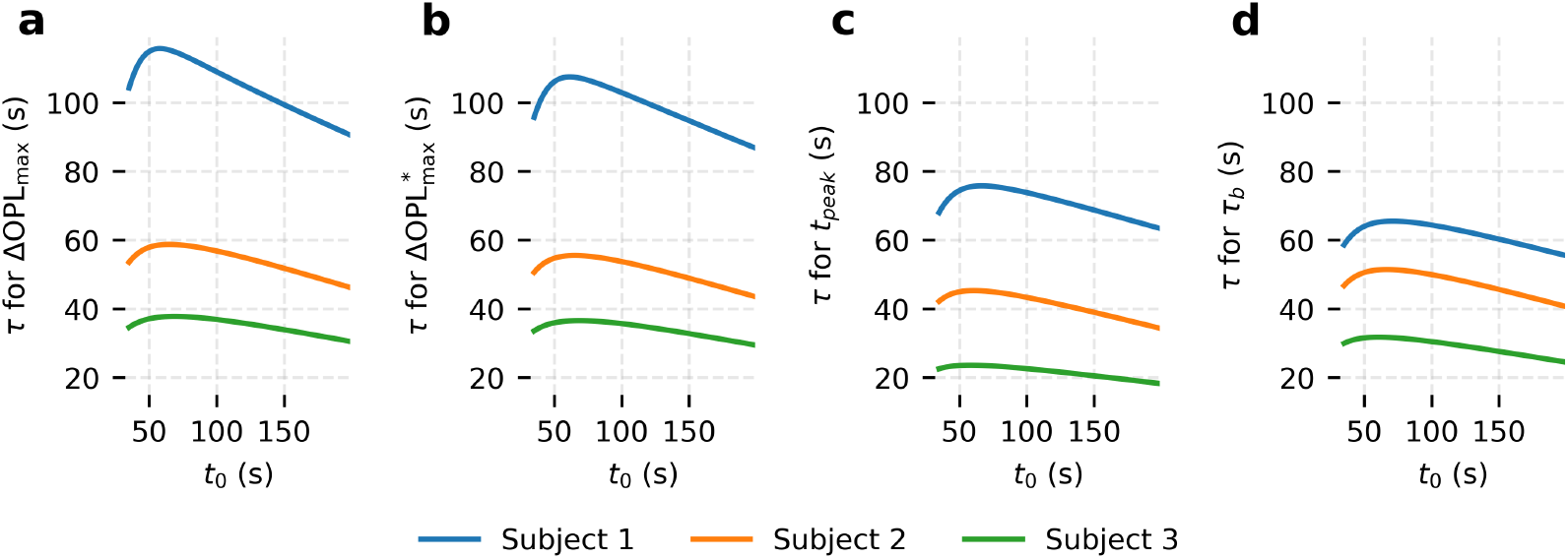
Dependence of the fitted adaptation time constant *τ* in Eq. 5 on the assumed cone pigment regeneration time constant *t*_0_ in Eq. A1. Panels show *τ* for four ORG metrics: **a** ΔOPL_max_, **b** 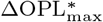, and **d** *τ*_*b*_, plotted as a function of *t*_0_ (30–200 s) for three subjects. For all metrics and subjects, the variation of *τ* with *t*_0_ remains within ~20%, and the relative ordering across subjects and metrics is preserved.

In the rate-limited model, the fraction of unbleached pigment *p*(*t*) is given by [25]:

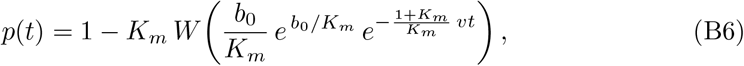

where *K*_*m*_ is the Michaelis constant, *W* is the Lambert–*W* function, *b*_0_ is the initial fractional bleach, and *v* is the initial rate of recovery following a total bleach. Here, *p*(*t*) represents the fraction of unbleached pigment available at time *t* after the initial high-bleach flash. To simulate pigment regeneration, We used *K*_*m*_ = 0.2 and *v* = 0.5 min^−1^ measured in [25], and set *b*_0_ = 0.67 as 67 % initial bleach.

Analogous to the procedure described in Eq. A3 - Eq. A5 for the exponential model, we firstly used the rate-limited pigment regeneration curve to compute a dark adaptation time, *t*_adapt_, for each ORG measurement. At each probe-flash time point, the available pigment fraction *p*(*t*) was evaluated using Eq. B6; this pigment level was then mapped back onto the unperturbed rate-limited recovery trajectory to determine the corresponding *t*_adapt_. ORG metrics aggregated across all trials in each subject were then expressed as a function of *t*_adapt_ and fit using a rate-limited scaling relation,

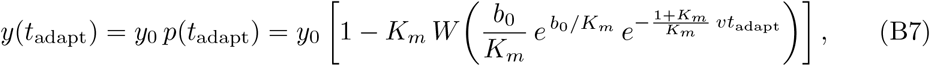

where *y*_0_ denotes the asymptotic dark-adapted ORG response amplitude. Under the assumption that the ORG metrics scale approximately with the fraction of unbleached pigment, Eq. B7 provides a direct link between pigment regeneration and the measured ORG responses.

In the fitting, *K*_*m*_ was held fixed at 0.2, consistent with prior work showing that the goodness of fit is only weakly dependent on *K*_*m*_ over the range 0.1–0.3 [25]. The remaining parameters *v, y*_0_, and *b*_0_ were treated as global fit parameters optimized across all trials. Figure B3 demonstrated the resulting rate-limited fits for three ORG metrics: (a-c) the amplitude ΔOPL_max_, (d-f) the time-to-peak *t*_pk_, and (h-j) the late contraction rate *τ*_*b*_.

To compare decay behavior across parameters and subjects within this framework, we used the parameter *v* as the primary descriptor of decay speed, in analogy to the time constants *τ* obtained from the exponential and probabilistic models (Methods 1 and 2). Across subjects, we observed the same qualitative trends as in those models. Subject 1 exhibited the slowest decay, with *v* values approximately one-half to one-third of those in subject 3 (0.22 min^−1^ versus 0.41 min^−1^ for ΔOPL_max_; 0.27 min^−1^ versus 0.71 min^−1^ for *t*_pk_; and 0.42 min^−1^ versus 0.80 min^−1^ for *τ*_*b*_). Furthermore, consistent with the exponential and probabilistic analyses, the ORG amplitude ΔOPL_max_ tended to recover more slowly (smaller *v*) than the temporal metrics *t*_pk_ and *τ*_*b*_, reinforcing the hypothesis that ORG amplitude and temporal dynamic might reflect partially distinct aspects of the cone recovery process.

We also assessed the fitting performance of the exponential and rate-limited models by comparing their coefficients of determination, *R*^2^. Across all subjects and metrics, the *R*^2^ values for the rate-limited fits (Fig. B3) were very similar to those obtained with the exponential fits (Figs. 4 and 5). In our dataset, therefore, the rate-limited model did not present a significant difference in goodness-of-fit over the exponential model. This is likely that rate-limited behavior is more accurate at lower or steady-state bleaching levels [15, 25], whereas our measurements followed a brief (35 ms) flash that produced a relatively high initial bleach (67 %).

Under these conditions, both models capture the main features of cone recovery, and the choice of model does not affect the qualitative conclusions about inter-subject variability or the relative behavior of amplitude versus temporal ORG metrics.

## Appendix C Method 3: Perturbed exponential model accounting for the flash-induced perturbations

To interpret the temporal evolution of ORG metrics during dark adaptation, we developed an exponential–step model combining the gradual exponential decay following the initial high photobleach with the perturbations caused by the probing flashes, which describes the decay of ORG parameters as a function of acquisition time *t*_acq_. It combines continuous exponential decay with multiplicative steps of magnitude *η* that represent the transient bleaching effects caused by the sequential probing flashes. The model proceeds iteratively through two steps for each measurement:

**Fig. B3.**
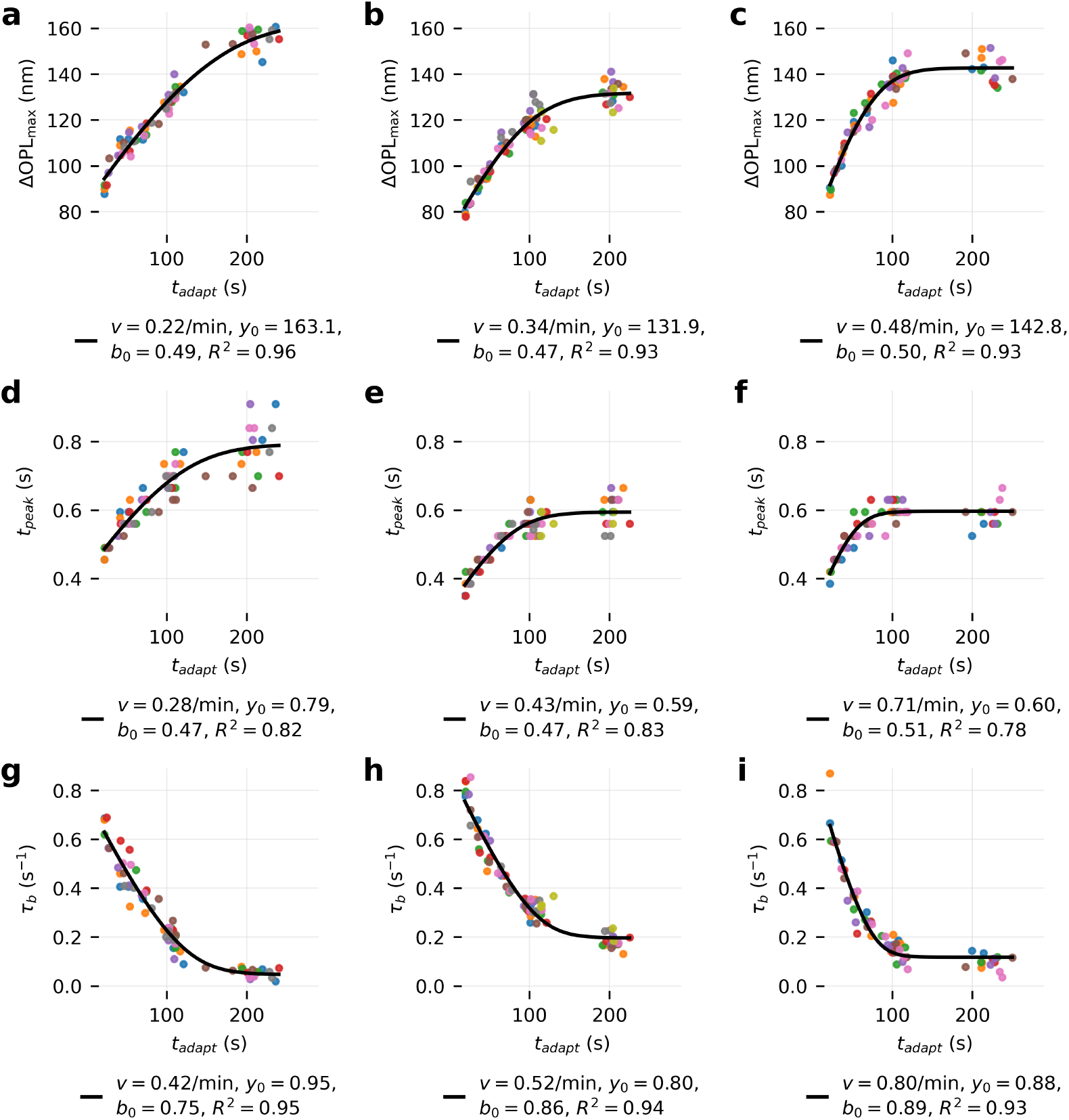
Rate-limited model fitting of ORG metrics (stimuli: 2.83 × 10^6^ photons*/*µm^2^): ΔOPLmax in **a, b, c**, *t*_pk_ in **d, e, f**, and *τ*_*b*_ in **g, h, i** as a function of adaptation time *t*_adapt_, computed from the rate-limited pigment regeneration model during cone dark adaptation, across 8-10 trials from subjects 1–3. Colored markers indicate fitted parameters from individual ORG measurements, with color indicating measurements acquired in the same dark adaptation trial. Solid black lines indicate rated-limited fits (Eq. B7, *Km* = 0.2) to the data, with resulting fitting parameters printed below each panel.

1. Perturbation induced by the probing flash at *t*_*i*−1_:

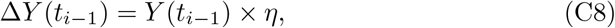

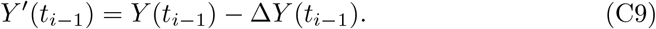
2. Exponential decay between two measurement times *t*_*i*−1_ *and t*_*i*_:

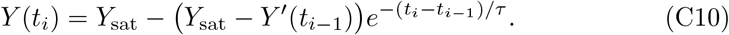

Iterating these two steps across all experimental acquisition times {*t*_*i*_} generates the complete fitted decay curve used to estimate the parameters *τ, η, Y*_sat_, and *Y*_0_.

Specifically, the ORG parameter *Y* (*t*_*i*_) evolves toward a saturation value *Y*_*sat*_ (the asymptotic dark-adapted limit), modeled by an exponential form with a time constant *τ* (see Eq. C10) for the recovery in dark. To account for perturbations induced by the test flash, a multiplicative step of magnitude *η* is applied at each acquisition time.

This hybrid exponential–step model effectively captures (1) the exponential decay of the ORG metrics quantified by the decay time constant *τ*, and (2) the perturbations (*η*) introduced by pigment bleach from repeated probing flashes during the dark adaptation.

Figure C4 shows an example of the probabilistic fitting applied to ΔOPL_max_ and *τ*_*b*_ from a single ORG measurement in subject 1. By iterating the two steps described above over all measurement times {*t*_*i*_} and minimizing the RMS fitting error, the optimized fitting curve is shown by the black line in Fig. C4. For ΔOPL_max_, the value increased from 71 nm to a saturation level of 165 nm, with a fitted decay time constant of 95 s. The test flash introduced a perturbation represented by *η* = 7 %, and the RMS fitting error was small at 2.37 nm. On the other hand, *τ*_*b*_ decreased from 0.76 s^−1^ to 0.03 s^−1^, with a time constant of 87 s and a perturbation of 13 %.

**Fig. C4.**
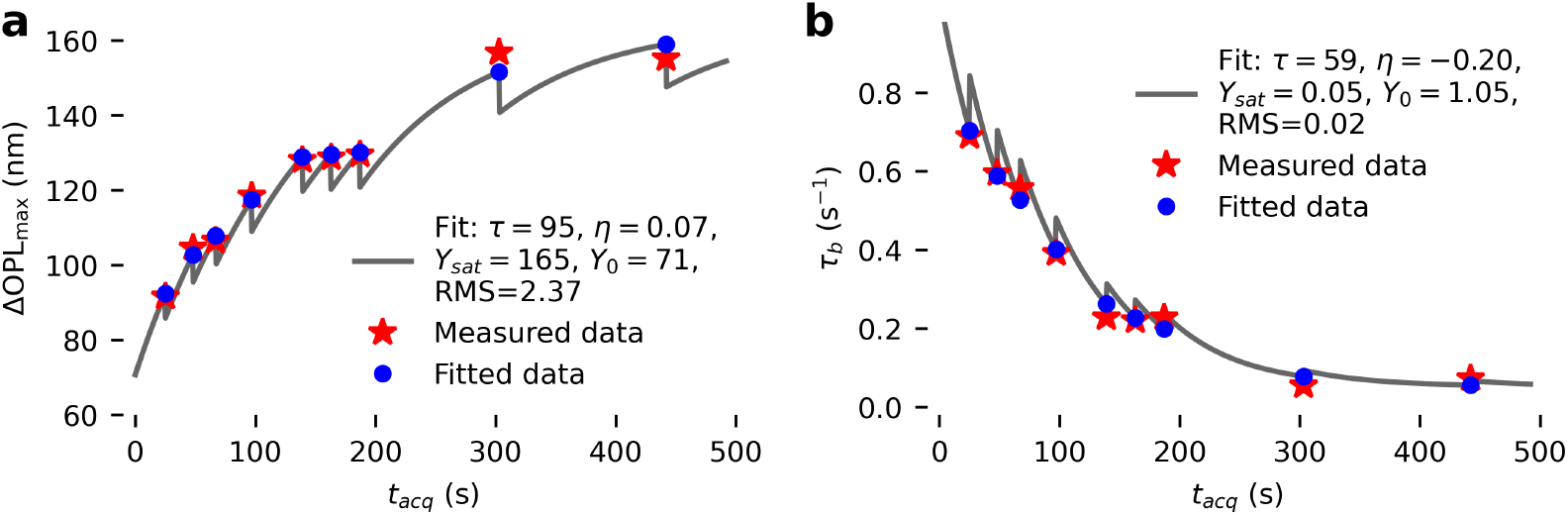
Probabilistic model fitting of cone ORG parameters. Representative fits of **a**, cone OS maximum OPL length change (ΔOPL_max_) and **b**, late contraction rate (*τ*_*b*_) as a function of acquisition time during dark adaptation (Subject 1, Trial 2). Measured data (red stars) are overlaid with the fitted results (blue circles). The solid curve denotes the probabilistic fitting model (Eq. C10) with the optimized model parameters (*τ, η, Y*_sat_, *Y*_0_) printed in the legend.

We then applied this model to all 8–10 trials collected from each subject. The global fitting parameters (*τ, η, Y*_sat_, *Y*_0_) were constrained to be common in all trials and optimized by minimizing the sum of squares error across the aggregated data. The resulting coefficients of determination (*R*^2^) were above 0.9 for both ΔOPL_max_ and *τ*_*b*_, and the values for *t*_pk_ ranged from 0.77 to 0.85, which are consistent with the *R*^2^ values obtained from Method 1 fitting shown in Fig. 4 and Fig. 5.

Additionally, the recovery time constants (*τ*) showed clear variations across subjects and ORG parameters (Table C1). These trends closely mirrored the results from Method 1 based on exponential pigment regeneration (Table 1). Specifically, the ORG amplitude recovered more slowly (average adaptation time constant of 71 s for ΔOPL_max_) than the parameters related to the ORG dynamics (average adaptation time constant of 55 s for *t*_pk_ and 58 s for *τ*_*b*_). Across subjects, the time constants (*τ*) decreased from Subject 1 to Subject 3.

These findings demonstrate the complementary interpretations offered by the other two fitting approaches. The perturbed exponential fitting relative to acquisition time provides a mathematical description of how the ORG parameters evolve with probing flashes during dark adaptation. This approach quantifies both their overall recovery and the variability introduced by the test-flash perturbations. Crucially, it achieves this without relying on fixed assumptions regarding the pigment regeneration time constant (*τ* = 60 s) used for the pigment level simulation in Method 1 (Appendix A) or the initial recovery rate (*v* = 0.5 min) used in Method 2 (Appendix B).

**Table C1.** Summary of the decay time constants for cone ORG parameters during dark adaptation using perturbed exponential model. The decay time constant *τ* (s) was derived through exponential-step fitting (Eq. C10 of cone response amplitudes (ΔOPL_max_) and temporal parameters (*t*_pk_ and *τ*_*b*_) as functions of acquisition time *t*_acq_.

| Subject | $\Delta OPL_{max}$ | $t_{pk}$ | $\tau_b$ |
| --- | --- | --- | --- |
| Subject 1 | 98 | 86 | 88 |
| Subject 2 | 62 | 48 | 55 |
| Subject 3 | 54 | 32 | 30 |
| Mean | 71 | 55 | 58 |

## References

[1] Lamb, T.D., Pugh, E.N.: Dark adaptation and the retinoid cycle of vision. Progress in Retinal and Eye Research 23(3), 307–380 (2004) 10.1016/j.preteyeres.2004.03.001

[2] Jiang, X., Mahroo, O.A.: Human retinal dark adaptation tracked in vivo with the electroretinogram: Insights into processes underlying recovery of cone-and rod-mediated vision. The Journal of Physiology 600(21), 4603–4621 (2022) 10.1113/JP283105

[3] Kefalov, V.J.: Rod and cone visual pigments and phototransduction through pharmacological, genetic, and physiological approaches*. Journal of Biological Chemistry 287(3), 1635–1641 (2012) 10.1074/jbc.R111.303008

[4] Nigalye, A.K., Hess, K., Pundlik, S.J., Jeffrey, B.G., Cukras, C.A., Husain, D.: Dark Adaptation and Its Role in Age-Related Macular Degeneration. Journal of Clinical Medicine 11(5) (2022) 10.3390/jcm11051358

[5] Weiss, E.: Shedding light on dark adaptation. The Biochemist 42(5), 44–50 (2020) 10.1042/BIO20200067

[6] Kalloniatis, M., Luu, C.: Light and Dark Adaptation. In: Kolb, H., Fernandez, E., Jones, B., Nelson, R. (eds.) Webvision: The Organization of the Retina and Visual System. University of Utah Health Sciences Center, Salt Lake City (UT) (1995)

[7] Pianta, M.J., Kalloniatis, M.: Characterisation of dark adaptation in human cone pathways: An application of the equivalent background hypothesis. The Journal of Physiology 528(Pt 3), 591–608 (2000) 10.1111/j.1469-7793.2000.00591.x

[8] Fain, G.L., Matthews, H.R., Cornwall, M.C., Koutalos, Y.: Adaptation in Vertebrate Photoreceptors. Physiological Reviews 81(1), 117–151 (2001) https://doi.org/10.1152/physrev.2001.81.1.117

[9] Kolesnikov, A.V., Kiser, P.D., Palczewski, K., Kefalov, V.J.: Function of Mammalian M-Cones Depends on the Level of CRALBP in Müller Cells. Journal of General Physiology 153(e202012675) (2020) 10.1085/jgp.202012675

[10] Saha, A., Bucci, T., Baudin, J., Sinha, R.: Regional tuning of photoreceptor adaptation in the primate retina. Nature Communications 15(1), 8821 (2024) 10.1038/s41467-024-53061-3

[11] Lamb, T.D., Pugh, E.N. Jr: Phototransduction, Dark Adaptation, and Rhodopsin Regeneration The Proctor Lecture. Investigative Ophthalmology & Visual Science 47(12), 5138–5152 (2006) 10.1167/iovs.06-0849

[12] Baumann, Ch.: Boll’s phenomenon. Vision Research 17(11), 1325–1327 (1977) 10.1016/0042-6989(77)90116-X

[13] Yang, G.-Q., Chen, T., Tao, Y., Zhang, Z.-M.: Recent advances in the dark adaptation investigations. International Journal of Ophthalmology 8(6), 1245– 1252 (2015) 10.3980/j.issn.2222-3959.2015.06.31

[14] Dimitrov, P.N., Guymer, R.H., Zele, A.J., Anderson, A.J., Vingrys, A.J.: Measuring Rod and Cone Dynamics in Age-Related Maculopathy. Investigative Ophthalmology & Visual Science 49(1), 55–65 (2008) 10.1167/iovs.06-1048

[15] Binns, A., Margrain, T.H.: Evaluation of retinal function using the Dynamic Focal Cone ERG. Ophthalmic and Physiological Optics 25(6), 492–500 (2005) 10.1111/j.1475-1313.2005.00338.x

[16] Owsley, C., Huisingh, C., Jackson, G.R., Curcio, C.A., Szalai, A.J., Dashti, N., Clark, M., Rookard, K., McCrory, M.A., Wright, T.T., Callahan, M.A., Kline, L.B., Witherspoon, C.D., McGwin, G. Jr: Associations Between Abnormal Rod-Mediated Dark Adaptation and Health and Functioning in Older Adults With Normal Macular Health. Investigative Ophthalmology & Visual Science 55(8), 4776–4789 (2014) 10.1167/iovs.14-14502

[17] Rushton, W.A.H., Henry, G.H.: Bleaching and regeneration of cone pigments in man. Vision Research 8(6), 617–631 (1968) 10.1016/0042-6989(68)90040-0

[18] Norren, D.V., Padmos, P.: Dark adaptation of separate cone systems studied with psychophysics and electroretinography. Vision Research 14(8), 677–686 (1974) 10.1016/0042-6989(74)90064-9

[19] Jackson, G.R., Scott, I.U., Kim, I.K., Quillen, D.A., Iannaccone, A., Edwards, J.G.: Diagnostic Sensitivity and Specificity of Dark Adaptometry for Detection of Age-Related Macular Degeneration. Investigative Ophthalmology & Visual Science 55(3), 1427–1431 (2014) 10.1167/iovs.13-13745

[20] Rushton, W.A.H.: Dark-adaptation and the regeneration of rhodopsin. The Journal of Physiology 156(1), 166–178 (1961) 10.1113/jphysiol.1961.sp006666

[21] Rushton, W.A.H.: Rhodopsin measurement and dark-adaptation in a subject deficient in cone vision. The Journal of Physiology 156(1), 193–205 (1961) 10.1113/jphysiol.1961.sp006668

[22] Geisler, W.S.: The effects of photopigment depletion on brightness and threshold. Vision Research 18(3), 269–278 (1978) 10.1016/0042-6989(78)90161-X

[23] Thomas, M.M., Lamb, T.D.: Light adaptation and dark adaptation of human rod photoreceptors measured from the a-wave of the electroretinogram. The Journal of Physiology 518(2), 479–496 (1999) 10.1111/j.1469-7793.1999.0479p.x

[24] Tillman, M.A., Panorgias, A., Werner, J.S.: Age-related change in fast adaptation mechanisms measured with the scotopic full-field ERG. Documenta Ophthalmologica 132(3), 201–212 (2016) 10.1007/s10633-016-9541-2

[25] Mahroo, OAR., Lamb, TD.: Recovery of the human photopic electroretinogram after bleaching exposures: Estimation of pigment regeneration kinetics. The Journal of Physiology 554(Pt 2), 417–437 (2004) 10.1113/jphysiol.2003.051250

[26] Mahroo, O.A.R., Lamb, T.D.: Slowed recovery of human photopic ERG a-wave amplitude following intense bleaches: A slowing of cone pigment regeneration? Documenta Ophthalmologica 125(2), 137–147 (2012) 10.1007/s10633-012-9344-z

[27] Kenkre, JS., Moran, NA., Lamb, TD., Mahroo, OAR.: Extremely rapid recovery of human cone circulating current at the extinction of bleaching exposures. The Journal of Physiology 567(Pt 1), 95–112 (2005) 10.1113/jphysiol.2005.088468

[28] Jonnal, R.S., Kocaoglu, O.P., Wang, Q., Lee, S., Miller, D.T.: Phase-sensitive imaging of the outer retina using optical coherence tomography and adaptive optics. Biomedical Optics Express 3(1), 104–124 (2012) 10.1364/BOE.3.000104

[29] Hillmann, D., Spahr, H., Pfäffle, C., Sudkamp, H., Franke, G., Hüttmann, G.: In Vivo Optical Imaging of Physiological Responses to Photostimulation in Human Photoreceptors. Proceedings of the National Academy of Sciences of the United States of America 113(46), 13138–13143 (2016) 10.1073/pnas.1606428113

[30] Azimipour, M., Migacz, J.V., Zawadzki, R.J., Werner, J.S., Jonnal, R.S.: Functional retinal imaging using adaptive optics swept-source OCT at 1.6 MHz. Optica 6(3), 300–303 (2019) 10.1364/OPTICA.6.000300

[31] Jonnal, R.S.: Toward a clinical optoretinogram: A review of noninvasive, optical tests of retinal neural function. Annals of Translational Medicine 9(15), 1270 (2021) 10.21037/atm-20-6440

[32] Chen, S., Huang, D., Jonnal, R.S.: Recent Developments in Photoreceptor Optoretinography and Progress Toward Clinical Use. Translational Vision Science & Technology 14(12), 18 (2025) 10.1167/tvst.14.12.18

[33] Zhang, P., Zawadzki, R.J., Goswami, M., Nguyen, P.T., Yarov-Yarovoy, V., Burns, M.E., Pugh, E.N.: In vivo optophysiology reveals that G-protein activation triggers osmotic swelling and increased light scattering of rod photoreceptors. Proceedings of the National Academy of Sciences 114(14), 2937–2946 (2017) 10.1073/pnas.1620572114

[34] Pandiyan, V.P., Maloney-Bertelli, A., Kuchenbecker, J.A., Boyle, K.C., Ling, T., Chen, Z.C., Park, B.H., Roorda, A., Palanker, D., Sabesan, R.: The Optoretinogram Reveals the Primary Steps of Phototransduction in the Living Human Eye. Science Advances 6(37), 1124 (2020) 10.1126/sciadv.abc1124. Chap. Research Article

[35] Pandiyan, V.P., Nguyen, P.T., Pugh, E.N., Sabesan, R.: Human cone elongation responses can be explained by photoactivated cone opsin and membrane swelling and osmotic response to phosphate produced by RGS9-catalyzed GTPase. Proceedings of the National Academy of Sciences 119(39), 2202485119 (2022) 10.1073/pnas.2202485119

[36] Tomczewski, S., Curatolo, A., Foik, A., Węgrzyn, P., Bałamut, B., Wielgo, M., Kulesza, W., Galińska, A., Kordecka, K., Gulati, S., Fernandes, H., Palczewski, K., Wojtkowski, M.: Photopic flicker optoretinography captures the light-driven length modulation of photoreceptors during phototransduction. Proceedings of the National Academy of Sciences 122(7), 2421722122 (2025) 10.1073/pnas.2421722122

[37] Zhang, F., Kurokawa, K., Lassoued, A., Crowell, J.A., Miller, D.T.: Cone photoreceptor classification in the living human eye from photostimulation-induced phase dynamics. Proceedings of the National Academy of Sciences 116(16), 7951–7956 (2019) 10.1073/pnas.1816360116

[38] Boyle, K.C., Chen, Z.C., Ling, T., Pandiyan, V.P., Kuchenbecker, J., Sabesan, R., Palanker, D.: Mechanisms of Light-Induced Deformations in Photoreceptors. Biophysical Journal 119(8), 1481–1488 (2020) 10.1016/j.bpj.2020.09.005

[39] Valente, D., Vienola, K.V., Zawadzki, R.J., Jonnal, R.S.: Insight into human photoreceptor function: Modeling optoretinographic responses to diverse stimuli. Science Advances 11(24), 7332 (2025) 10.1126/sciadv.adq7332

[40] Jackson, G.R., Owsley, C., McGwin, G.: Aging and Dark Adaptation. Vision Research 39(23), 3975–3982 (1999) 10.1016/s0042-6989(99)00092-9

[41] Ding, J., Kim, T.-H., Ma, G., Yao, X.: Intrinsic signal optoretinography of dark adaptation abnormality due to rod photoreceptor degeneration. Experimental Biology and Medicine (Maywood, N.J.) 249, 10024 (2024) 10.3389/ebm.2024.10024

[42] Zhang, P., Wahl, D.J., Mocci, J., Miller, E.B., Bonora, S., Sarunic, M.V., Zawadzki, R.J.: Adaptive optics scanning laser ophthalmoscopy and optical coherence tomography (AO-SLO-OCT) system for in vivo mouse retina imaging. Biomedical Optics Express 14(1), 299–314 (2023) 10.1364/BOE.473447

[43] Zhang, P., Karlen, S.J., Allina, G.P., Zawadzki, R.J.: Whole retina light-evoked optoretinography (ORG) under different retina hydration levels: Modeling of Bruch’s membrane ORGs. Biomedical Optics Express 16(5), 1944–1959 (2025) 10.1364/BOE.551862

[44] Wong, J.H., Luo, S., Hosseinaee, Z., Feroldi, F., Roorda, A.: Optoretinography with actively stabilized adaptive optics optical coherence tomography. Biomedical Optics Express 16(8), 3222–3236 (2025) 10.1364/BOE.566376

[45] Azimipour, M., Valente, D., Vienola, K.V., Werner, J.S., Zawadzki, R.J., Zawadzki, R.J., Jonnal, R.S.: Optoretinogram: Optical Measurement of Human Cone and Rod Photoreceptor Responses to Light. Optics Letters 45(17), 4658–4661 (2020) 10.1364/OL.398868

[46] Li, H., Weiss, C.E., Pandiyan, V.P., Nanni, D., Liu, T., Kung, P.W., Tan, B., Barathi, V.A., Schmetterer, L., Sabesan, R., Ling, T.: Optoretinography Reveals Rapid Rod Photoreceptor Movement upon Photoisomerization. bioRxiv (2025). 10.1101/2025.03.22.644466

[47] Azimipour, M., Jonnal, R.S., Werner, J.S., Zawadzki, R.J., Zawadzki, R.J.: Coextensive synchronized SLO-OCT with adaptive optics for human retinal imaging. Optics Letters 44(17), 4219–4222 (2019) 10.1364/OL.44.004219

[48] Jonnal, R.S.: Rjonnal/Ciao: Initial Release. Zenodo (2020). 10.5281/zenodo.3903941

[49] Stockman, A., Sharpe, L.T., Fach, C.: The spectral sensitivity of the human short-wavelength sensitive cones derived from thresholds and color matches. Vision Research 39(17), 2901–2927 (1999) 10.1016/S0042-6989(98)00225-9

[50] Otake, S., Gowdy, P.D., Cicerone, C.M.: The spatial arrangement of L and M cones in the peripheral human retina. Vision Research 40(6), 677–693 (2000) 10.1016/s0042-6989(99)00202-3

[51] Aggarwala, K.R., Kruger, E.S., Mathews, S., Kruger, P.B.: Spectral bandwidth and ocular accommodation. JOSA A 12(3), 450–455 (1995) 10.1364/JOSAA.12.000450

[52] Vienola, K.V., Valente, D., Zawadzki, R.J., Jonnal, R.S.: Velocity-based optoretinography for clinical applications. Optica 9(10), 1100–1108 (2022) 10.1364/OPTICA.460835

[53] Maddipatla, R., Cai, Y., Zawadzki, R.J., Jonnal, R.S.: Photopigment Bleaching Calculation and Simulation Notebook. Zenodo (2026). 10.5281/zenodo.18988166

[54] ANSI: American National Standard for Safe Use of Lasers Z136. 1-2014. Laser Institute of America Orlando, FL (2014)

[55] Cai, Y., Bartuzel, M.M., Maddipatla, R., Zawadzki, R.J., Jonnal, R.S.: Phase-based optoretinographic measurements of cones with a raster-scanning adaptive optics OCT are highly repeatable. Biomedical Optics Express 17(5), 2598–2609 (2026) 10.1364/BOE.591181

[56] Cai, Y.: Codebase and datasets for cone optoretinogram (ORG) dark adaptation analysis. Zenodo (2026). 10.5281/zenodo.19834538

